# Gene expression plasticity, genetic variation and fatty acid remodelling in divergent populations of a tropical bivalve species

**DOI:** 10.1101/2021.02.08.429829

**Authors:** J. Le Luyer, L. Milhade, C. Reisser, C. Soyez, C. J. Monaco, C. Belliard, G. Le Moullac, C.-L. Ky, F. Pernet

## Abstract

Ocean warming is a particularly challenging threat for tropical marine bivalves’ species because many live already near their upper thermal limits. The thermal sensitivity of organisms is a strong contributor to the biogeographic boundaries of populations and species. The potential of thermal plastic response (range of thermal breadth) is typically reduced for marine populations living in stable thermal environments, e.g., in extreme latitudes organisms as traduced in the climatic variability hypothesis. However, regional-scale heterogeneity among tropical environments, such as archipelagos in French Polynesia, might also serve in modulating this plastic potential. The questions remain now, how tropical organisms are able to cope with abnormally elevated temperature on long-term (several weeks) and how environmental-variability might drive the potential of resilience? To answer these questions, we benefit from two ecologically divergent populations of a marine tropical mollusc species, *Pinctada margaritifera*, that usually experience either large diurnal variations (tide-pools, Marquesas archipelago) or lower temperature with stable to moderate variations (Gambier archipelago). Individuals were maintained in common garden experiment at several controlled temperature conditions (23°C, 28°C, 32°C and 34°C) over a 48 days period. We explored genetic divergence as well as thermal plastic responses by combining lipidomic and transcriptomic approaches. We show that *P. margaritifera* have capacities to adjust to long-term elevated temperatures that was thus far largely underestimated. Furthermore, we identified genetic variation between populations that overlapped with genes expression variations, including genes involved in the respiration machinery, a central process delimiting critical temperatures in marine invertebrates. This study is the first of a series looking at the global adaptation and acclimation mechanisms in response to climate change in *Pinctada* species.

## Introduction

Temperature is a major determinant of marine species distribution (Hochachka & Somero, 2002). The expected increase in temperature caused by global warming is particularly challenging for stenothermal species and for those that already experiencing temperatures close to their upper thermal limit (Dahlke, Wohlrab, Butzin, & Pörtner, 2020; Tewksbury, Huey, & Deutsch, 2008). In the tropical black-lip pearl oyster (*Pinctada margaritifera*), the physiological thermal performance peaks at *ca*. 28-29°C, while at 22°C and 34°C growth and reproduction are suspended (Le Moullac et al., 2016). The seawater temperature in French Polynesian archipelagoes is usually below 31°C, with limited daily and annual variation (Van Wynsberge et al., 2017). However, temperatures recorded across the region reveal that in some lagoons *P. margaritifera* experiences temperatures warmer than their thermal optimum (28-29°C) during ~121 days per year. These populations therefore live close to their upper physiological limits (Le Moullac et al., 2016), a threat that is now being amplified by ongoing climate change. The contrasting thermal regimes that exist across French Polynesian lagoons offers an opportunity to explore mechanisms of thermal adaptation. Particularly interesting is the divergent intertidal population of the Marquesas archipelago, which experience daily temperatures ranging between 27°C and more than 34°C (Reisser et al., 2019). How these organisms perform under heat stress and which potential physiological capacities allow this population to cope with high and fluctuating temperatures remain unknown. We know however that Marquesas populations are genetically and phenotypically distinct from *P. margaritifera* populations present in other archipelagos, reflecting possible local adaptation and/or acclimation potential (i.e., phenotypic plasticity) (Reisser et al., 2019).

Phenotypic plasticity is generally viewed as a mechanism allowing individuals to survive to stochastic and/or highly variable environmental conditions (Fox, Donelson, Schunter, Ravasi, & Gaitán-Espitia, 2019; Seebacher, White, & Franklin, 2015). Plastic responses to temperature involve a wide-range of biological processes, including alterations of the velocity of chemical and enzymatic reactions, chaperones production, rates of diffusion, membrane fluidity and protein structure (Hochachka & Somero, 2002). Among these processes, membrane remodeling is pivotal. Increasing temperature increases membrane fluidity, which can lead to membrane dysfunction. Poikilotherms are able to counteract the effect of temperature on membrane structure by remodeling membrane lipids, *via* changes in phospholipid head groups, fatty acid composition and cholesterol content, a process known as homeoviscous adaptation (HVA) (Hazel, 1995). In bivalves, HVA can be accomplished by the decreased unsaturation of phospholipid acyl chains with increasing temperature (Hall, Parrish, & Thompson, 2002; Pernet, Tremblay, Comeau, & Guderley, 2007). For example, in temperate bivalve species (e.g., the scallop *Placopecten magellanicus*, the oysters *Crassostrea gigas* and *C. virginica*, and the blue mussel, *Mytilus edulis*), the long-chain polyunsaturated fatty acids increase with decreasing acclimation/acclimatization temperature (Delisle et al., 2020; Hall et al., 2002; Pernet et al., 2007). Data remain scarce for tropical species, yet the giant clam *Tridacna maxima* shows membrane lipid remodeling in response to temperature consistent with HVA (Dubousquet et al., 2016).

Genome-wide transcriptomic approaches can uncover additional details on the repertoire of molecular and cellular mechanisms responsible for thermal plasticity (Evans, 2015; Lee, Taylor, Shen, & Ehrenreich, 2016; Yeaman, 2015). For example, transcriptomics and proteomics have revealed the complex interplay between chaperone regulation, redox and metabolic shifts exhibited by thermally-stressed bivalves (Jiawei Chen, Liu, Cai, & Zhang, 2019; Tomanek, 2010). The potential for acclimation to thermal stress also depends on the amplitude and/or frequency of the environmental current and past variation (Donelson, Munday, McCormick, & Pitcher, 2011; Donelson, Salinas, Munday, & Shama, 2018;

Kellermann, van Heerwaarden, & Sgrò, 2017). According to the climatic variability hypothesis (CHV), the potential for thermal plasticity is higher for species or populations inhabiting highly-variable environments (e.g., temperate latitudes) than for those that routinely experience little change (e.g., extreme low and high latitudes) (Bozinovic, Calosi, & Spicer, 2011; Bozinovic & Pörtner, 2015; Dahlke et al., 2020). Finally, the potential for thermal plasticity is also largely dependent on prior acclimation to stressful conditions (Dahlke et al., 2020; Seebacher et al., 2015) and the duration of the stress (Semsar-kazerouni & Verberk, 2018), with inter-individual variability in response that can persist across generations, i.e. transgenerational plasticity (Donelson et al., 2011; Donelson, Wong, Booth, & Munday, 2016).

Environmental pressure might also favor the frequency of adaptive alleles at the scale of the population which would confer heritable changes modifications to specific environmental conditions. For instance, natural selection has contributed to the higher thermal tolerance and increased invasive potential of the blue *M. galloprovencialis* compared to other *Mytilus* species (Popovic & Riginos, 2020; Saarman, Kober, Simison, & Pogson, 2017). In *C. gigas*, natural selection and/or local adaption on thermal performances shaped the fine-scale structure of populations living in a highly-connected and temperature-contrasted area (Li et al., 2018). The genetic divergence observed among these populations of *C. gigas* also correlates with phenotypic divergence in the response to thermal stress, and suggests that plasticity might be favorably selected for populations subject to more variable environments (Li et al., 2018). Efforts to quantify the contributions of genetics and plastic responses to temperature exhibited by organisms are challenging, but critical for implementing sound conservation policies and ensuring the persistence of natural and managed populations under climate change (Chevin, Lande, & Mace, 2010; Duarte et al., 2020). In the Pacific basin, local adaptation associated with heterogenic environmental conditions was suggested to explain the weak structure of the *P. margaritifera* population in five main clusters (Lal, Southgate, Jerry, Bosserelle, & Zenger, 2017). In French Polynesia, the *P. margaritifera* populations from Marquesas and Gambier archipelagoes live under contrasting environments and exhibit strong genetic differentiation (Reisser et al., 2019); however, no study has explored the possible adaptive potential underlying this structuring.

Here we examine the thermal sensitivity of *P. margaritifera* by combining complementary transcriptomic and lipidomic approaches among populations. Using two ecologically divergent populations (Gambier population: stable – cold *vs*. Marquesas tidepool population: highly variable – hot environment), we quantify the adaptive *vs*. plastic response to temperatures over different timescales (from days to several weeks).

## Material and methods

Marquesan breeders were collected from tidepools (Ua Pou; French Polynesia; 9°25′18.7″S, 140°03′07.5″W) and air-shipped to Ifremer facilities (Taravao, Tahiti, French Polynesia; 17°48′31.7″S, 149°17′41.4″W). Five females were crossed with seven males under a full-factorial breeding scheme according to a standard procedure developed in the laboratory (Ky, Sham Koua, & Le Moullac, 2018). Concurrently, in a local private hatchery at Gambier (CA Regahiga Pearls Farm & Hatchery; Mangareva, French Polynesia; 23°07′05.2″S, 134°58′56.8″W), eight females were crossed with ten males (full-factorial breeding scheme). Progenies from the Marquesas and Gambier crosses were then transferred to a lagoon nearby Ifremer (Tahiti), and maintained for one month before the acclimation period in the laboratory (September 2017). At the start of the experiment, 16 individuals per population were sampled randomly to examine the initial physiological condition. Animals’ cohorts were of comparable conditions with a mean wet weight (g) of 30.80 ± 6.96 and 35.04 ± 8.74 (se), for Gambier and Marquesas, respectively (t-test; t = −1.53; *P* = 0.14) and mean shell width (cm) of 6.60 ± 0.60 and 6.70 ± 0.58 (se) for Gambier and Marquesas, respectively (t-test; t = −0.51; *P* = 0.61).

Individuals from Marquesas were slightly longer than those from Gambier, with a mean length (cm) of 6.39 ± 0.49 and 6.82 ± 0.46 (se), for Gambier and Marquesas, respectively (t-test; t = −2.56; *P* = 0.02). Individuals sampled at the start of the experiment were all males.

### (a) Experimental design, physiological measurements, and tissue sampling

After the one-month acclimation in the lagoon, animals were further acclimated for three weeks to the laboratory conditions at 27°C (*i.e*. ambient). A total of 96 individuals from each population were placed in flow-through raceways (16 oysters per 20-L tanks nested in 500-L raceways per temperature condition). Animals were supplied continuously (*ad libitum*) with a mix of *Isochrysis galbana* and *Chaetoceros minus* microalgae, at a daily ratio equal to 7–8% dry-weight-algae/dry-weight-oyster, for limiting pseudo-faeces production (Gardon, Reisser, Soyez, Quillien, & Le Moullac, 2018). Temperatures were recorded hourly using iBWetLand biochip (Alpha Mach inc., Canada) throughout the experiment. Seawater was heated by an electric heater or cooled with a heat exchanger (calorie exchange with cold freshwater), both operated by a temperature controller.

At the end of the acclimation period (October 6), animals were subjected to a gradual decrease/increase in temperature, reaching either 23°C (cooling), 32°C or 34°C (warming) within 2 days. Another group was kept at 27°C. These temperatures reflect normal/reference (27°C), extreme-cold (23°C), warm (32°C) and extreme-hot (34°C) conditions for oysters in French Polynesia. Note that the temperature treatments were not replicated due to logistic constraint, thus precluding our ability to unambiguously separate the treatment effect from among-tank differences (Hurlbert, 1984).

Animals were maintained at their respective temperatures for 48 days. Individuals were sampled on days 2 and 48 for molecular (transcriptomics) analyses, on days 48 for lipids and on days 0, 2, 28 and 48 for biometry (wet weight, length and sex determination). Each sampling consisted in four oysters per combination of population x temperature x time. Sex was determined by microscopic examination of the gametes. Two pieces of mantle tissue from the same area (approx. 1 cm^2^) were stored in RNA-Later at −80°C or in chloroform/methanol (2:1, v/v) at −20°C for molecular and lipid analysis, respectively. The sampling was done at the same time of the day across all sampling dates to prevent confounding circadian-rhythm effects with genes expression (Doherty & Kay, 2010).

### (b) Lipids

Sample handling, storage and analysis followed the recommendations for best practices for lipid research in aquatic sciences (Couturier et al., 2020). Briefly, tissue samples were homogenized over crushed ice using a glass pestle and a 7-mL Tenbroeck tissue grinder containing 2 mL of chloroform/methanol (2:1, v/v). The solution was poured in a 6-mL glass vial and rinsed twice with 1-mL chloroform/methanol (2:1, v/v). Samples were capped with Teflon and kept in nitrogen at −20°C f and analysed within one month.

To study the fatty acid remodeling of membrane lipids in response to temperature, it is often necessary to purify polar lipids (Couturier et al., 2020). Indeed, the neutral lipids, which constitute energy reserves, are not involved in this process but may contain fatty acids. Here we analyzed neutral lipids by thin layer chromatography and found only sterols (no free fatty acids, alcohols, mono-diacylglycerols, triacylglycerols). We therefore analyze the fatty acid composition of bulk lipids (no purification), considering that they come exclusively from polar lipids (Couturier et al., 2020).

A 1-mL subsample of the chloroform/methanol (2:1, v/v) mixture was evaporated to dryness under nitrogen. A known amount of 23:0 fatty acid was added as an internal standard. Fatty acid methyl esters (FAME) were obtained by acidic transesterification. Briefly, 0.8 mL of H_2_SO_4_/methanol was added (3.4%, v/v) and lipid samples were then vortexed and heated at 100°C for 10 min. After cooling, 0.8 mL of hexane and 1.5 mL of distilled water saturated in hexane were added. Samples were homogenized and centrifuged at 738 g for 1 min at room temperature. The aqueous phase (without FAME) was discarded and the organic phase– containing FAME was washed twice with distilled water saturated in hexane. This transesterification produces fatty acid methyl esters (FAME) from the fatty acid esterified at the sn-1 and sn-2 position of diacylphospholipids, and the sn-2 position of plasmalogen phospholipids. It also produces dimethyl acetals (DMA) from the alkenyl chains at the sn-1 position of plasmalogens (Morrison & Smith, n.d.). FAME and DMA were analysed in a HP6890 gas-chromatography system (Hewlett-Packard) equipped with a DB-Wax capillary column (30 m × 0.25 mm; 0.25-μm film thickness; Agilent technologies) and a flame ionization detector. Hydrogen was used as the carrier gas. The fatty acids were identified by comparing their retention times with those of standards (Supelco 37 Component FAME Mix, the PUFA No.1 and No.3, and the Bacterial Acid Methyl Ester Mix from Sigma) and in-house standard mixtures from marine bivalves and microalgae. Fatty acid compositions were expressed as the mass percentage of the total fatty acid content. Here we particularly focused on polyunsaturated fatty acids (PUFAs) and unsaturation index, *i.e*. the average number of double bonds per acyl chain, because they generally vary with temperature in a way consistent with HVA in other bivalves (Hazel, 1995; Pernet et al., 2007).

### (c) Statistics for biometry and fatty acids

Statistics were done using R (R Core Team, 2012) and differences were considered significant when *P* < 0.05, unless otherwise specified.

#### Biometry

Variation in weight data was assessed for each time point using ANOVA with population (two levels), temperature (four levels) and time (three levels) as explanatory variables. Normality and homoscedasticity were assessed visually and statistically (Shapiro and Levene tests, respectively). When significant effects were detected, Tukey’s HSD post-hoc tests were used to assess variation for factors with more than two levels. Variation in sex categories was assessed using the Chi-squared test. When significant effects were detected, we computed the contribution (%) of each sex category to the difference observed (calculated as follow: contribution (%) = r^2^ / χ^2^, with *r* the Pearson residual for each cell (standardized residual) and *χ* the Chi-square statistic.

#### Fatty acids

We used a subset of the most common fatty acid type (at least 1% of the total FA content) to assess the variation explained by both temperature and population on the FA content on day 48. We first computed a Euclidian distance matrix on the FA content across individuals and performed a principal coordinate analysis (PCoA) on this Euclidian distance matrix. Only PCo factors showing a relative eigenvalue higher than 2% (n = 6) were retained for the analysis, accounting for 97.3% of the total variance. The Euclidean distance and the PCoA were computed, respectively, using the functions ‘*daisy*’ and ‘*pcoa*’ available in the *ape* R package (Paradis, Claude, & Strimmer, 2004). We first produced a stepwise model selection on variables temperature and population using the function ‘*ordistep*’ in the *vegan* R package (Oksanen et al., 2012). A distance-based redundancy analysis (db-RDA) was then computed using the retained PCo factors as a response matrix and the variables temperature and population as the explanatory factors. Partial db-RDAs were produced to test for the effect of rearing temperature or population alone, while controlling for the other variable.

We used two-way ANOVA with interaction to assess the effects of population and temperature on specific FA on day 48. Pearson’s correlations were used to visualize pairwise similarities across FA classes; associations were considered significant when *P* < 0.01.

### (d) DNA/RNA extraction and sequencing

Total RNA was extracted from *P. margaritifera* by lacerating the mantle tissues with a scalpel and rinsing with 1X PBS. Cellular lysis was induced with 1.5 mL of TRIzol (Invitrogen, USA) according to the manufacturer’s recommendations. The supernatant was transferred into a 2-mL tube and incubated for 10 min on ice. The phase separation was achieved by adding 300 μL of chloroform followed by centrifugation at 12,000 x *g* for 12 min at 4°C. The upper aqueous layer contained the RNA, and the lower organic layer was stored at −20°C for later DNA extraction. Total RNA from each individual was subjected to a DNAse treatment using Qiagen’s RNA cleanup kit (USA). RNA and DNA were quantified using a NanoDrop ND-2000 spectrophotometer (Thermo-Fisher, USA), and RNA quality was further evaluated using a Bioanalyzer 2100 (Agilent, USA). High-quality RNA was sent to McGill University’s “Genome Quebec Innovation Center” (Montréal, QC, Canada) for Nextera XT (Illumina, USA) library preparation and sequencing on an Illumina HiSeq4000 100 bp paired-end platform, multiplexing 12 samples per lane.

### (e) Mapping and coding-region genotyping

Raw reads provided by RNA sequencing were filtered for quality and length using Trimmomatic v.0.36 (Bolger, Lohse, & Usadel, 2014) with the quality parameters minimum length, trailing, and leading set to 60 bp, 20, and 20, respectively. Illumina’s adaptors and residual cloning vectors were removed using the UNIVEC database (https://www.ncbi.nlm.nih.gov/tools/vecscreen/univec/). Trimmed paired-end reads were mapped against *P. margaritifera* reference genome (Le Luyer et al., 2019) using GSNAP v2018-07-04 (Wu, Reeder, Lawrence, Becker, & Brauer, 2016; --max-mismatches = 2; --novel-splicing =1; --min-coverage=0.95; --split-output). Only the uniquely-mapped reads (concordant_uniq) with mapping value > 5 were retained for downstream analyses.

For variants discovery (Single Nucleotide Polymorphisms; SNPs), mappings were processed with GATKv-4.0.3.0 suite (Auwera et al., 2013) following recommendations for RNAseq data. Raw SAM files were trimmed using ‘*CleanSam*’ and ‘*MarkDuplicates*’ functions to remove putative PCR duplicates (Auwera et al., 2013). The function ‘*SplitNCigarReads*’ was used to filter problematic reads according to their CIGAR string values (-refactor-cigar-string true; -RF CigarContainsNoNOperator; -RF GoodCigarReadFilter). A raw VCF file was produced using Freebayes-parallel, with default parameters except for the following: --use-best-n-alleles option set to 4, a minimum mapping quality –min-mapping-quality of 30, and a minimum coverage of 10 reads present across all samples (--min-coverage) for the locus. The VCFlib (https://github.com/vcflib/vcflib) program “vcffilter” was used to replace any population called with less than 10 reads for a given individual (-g “DP > 10”), keeping only SNP variants (-f “TYPE=SNP”). The resulting VCF was then filtered with vcftools (Danecek et al., 2011) to remove loci with a minimum allele frequency under 0.1, and more than 10% missing data (global). We also removed individuals bearing more than 15% missing data (4 individuals in total). The samtools distributed BCFtools (H. Li et al., 2009) was used to keep only biallelic sites (-m2 −M2). Finally, missing genotypes were imputed using a Hidden Markov Model approach implemented in Beagle v5.0 (Browning, Zhou, & Browning, 2018).

### (f) Genetic structuring, differentiation and candidate SNPs

The *vcfR* package (Knaus & Grünwald, 2016) was used to integrate the dataset into R. We did two exploratory analyses: first, we performed a Principal Component Analysis (PCA) using the *adegenet* package (Jombart & Ahmed, 2011) and the “*glPca*” function, with 3 PC axes kept, explaining 25.06, 4.68 and 3.91% of the variance, for PC1, PC2 and PC3, respectively. Then, we searched for the presence of genetic structuring using a non-supervised hierarchical clustering method with the “*find.clusters*” function implemented in the *adegenet* R package, and we kept 50 PC axes for analysis.

To search for putative candidate SNPs that significantly contribute to the differentiation of genetic clusters, we performed a Discriminant Analysis of Principal Coordinates (DAPC) as implemented in *adegenet*. To avoid overfitting the data by keeping too many PC axes, we used the K clusters identified in the exploratory step above, and performed a stratified cross-validation of DAPC with the function ‘*xvalDapc*’ in *adegenet* (the ‘best’ number of PC axes was 30). We then performed the DAPC retaining 78.04% of the total variance and extracted the loading contribution of each SNP to the global variance of the sample set. SNPs were then ranked according to their contribution and were defined as putative candidates if they were found in the top 1 percentile of the distribution (maximum contribution to the total variance).

We used *snpEff* (Cingolani et al., 2012) to assess the effect of candidate SNPs on the genes present in the genome of *P. margaritifera*. To test for potential gene ontology (GO) enrichment of the candidate SNPs that could identify important biological processes differing between the two populations, GO terms associated with transcripts carrying candidate SNPs were used in a Fisher’s test using the GOATOOLS python library (Klopfenstein et al., 2018) implemented in “go_enrichment” GitHub repository (https://github.com/enormandeau/go_enrichment). We retained enriched GO if *P* < 0.001. Finally, population genetic differentiation was estimated by computing the pairwise Fst between the populations with the R package *hierfstat* (Goudet, 2005).

### (g) Variance partitioning analysis of gene expression

For gene expression quantification, raw counts were computed using HTSeq v0.9.1 software (Anders, Pyl, & Huber, 2015) based on a custom-made GFF3 file [Supplementary Material (Le Luyer et al., 2019)]. Genes showing residuals expression (< 1 CPM in at least 4 individuals) were removed for downstream analysis using *DESeq2* v1.22.1 R package (Love, Huber, & Anders, 2014). To assess the relative contribution of population, time, and/or temperature on the gene expression of *P. margaritifera*, we followed the same procedure than for the lipids using combination of RDAs. For gene expression, the selection of the optimal number of PCoA axes was done using the broken-stick approach (Legendre & Gallagher, 2001; Legendre & legendre, 2012), which resulted in a total of 18 axes explaining 64.95% of the total variance. Three partial db-RDAs and analysis of variance (ANOVA, 1000 permutations) were used to validate that effect of each factor alone while controlling for the other factor. The effect of a given factor was considered significant when *P* values were < 0.05.

### (h) Differential gene expression and reaction norms classification

We explored the variability in gene expression across temperatures, populations and their interactions for each sampling time separately using a series of Generalized Linear Models (GLM) and Likelihood-ratio tests (LRT) implemented in *DESeq2* v1.22.1 R package (Love et al., 2014). We further explored variation in gene expression using pairwise group comparisons (per time x population x temperature) using contrasts and Wald tests (Yeaman et al., 2014). For contrasts, genes were considered differentially expressed if false discovery rate (FDR) < 0.05 and absolute fold-change (|FC|) was > 2. GO enrichment analyses were conducted following same procedure than described above and enrichment were considered significant for Bonferroni adj. *P* < 0.1.

We then used GLMs to identify genes with significant interaction (population x temperature). The full model containing the interaction term was retained if LRT *P* < 0.05. Finally, to classify the reaction norms of the temperature-responsive genes, we compared linear (temperature) and quadratic (temperature + temperature^2^) models for each time and population separately (Supplementary Methods and Table S2). When the quadratic model best fit the data (LRT adj. *P* < 0.05), we computed the optimal temperature (*Tm*) following previous published formula (Jun Chen, Nolte, & Schlötterer, 2015): *e = Em + g2(t – Tm)^2^*; with expression vector (*e*), the expression at *Tm* in quadratic equation (*Em*), the quadratic coefficient (*g2*) and the temperature (*t*).

## Results

### (a) Individual performance

Mortality was negligible throughout the experiment (< 2 %) and independent of the acclimation temperature or origin. Gametes were detected in oysters on all sampling dates and temperatures, but 75% of individuals had no gametes after 28 d at 34°C (Fig. 1). The absence of gametes in 34°C on day 28 mainly contributes (40.40%) to significantly differentiate temperature conditions (Pearson’s Chi-square; χ^2^ = 19.8; *P* = 0.02). The wet weight of animals was unaffected by temperature or population, but varied across time. Individuals sampled on day 48 were heavier than those sampled earlier (Tukey’s HSD; *P* = 0.02 and 0.04 for the 48d-2d and 48d-28d contrasts, respectively; Supplementary Fig. S1).

**Fig. 1:**
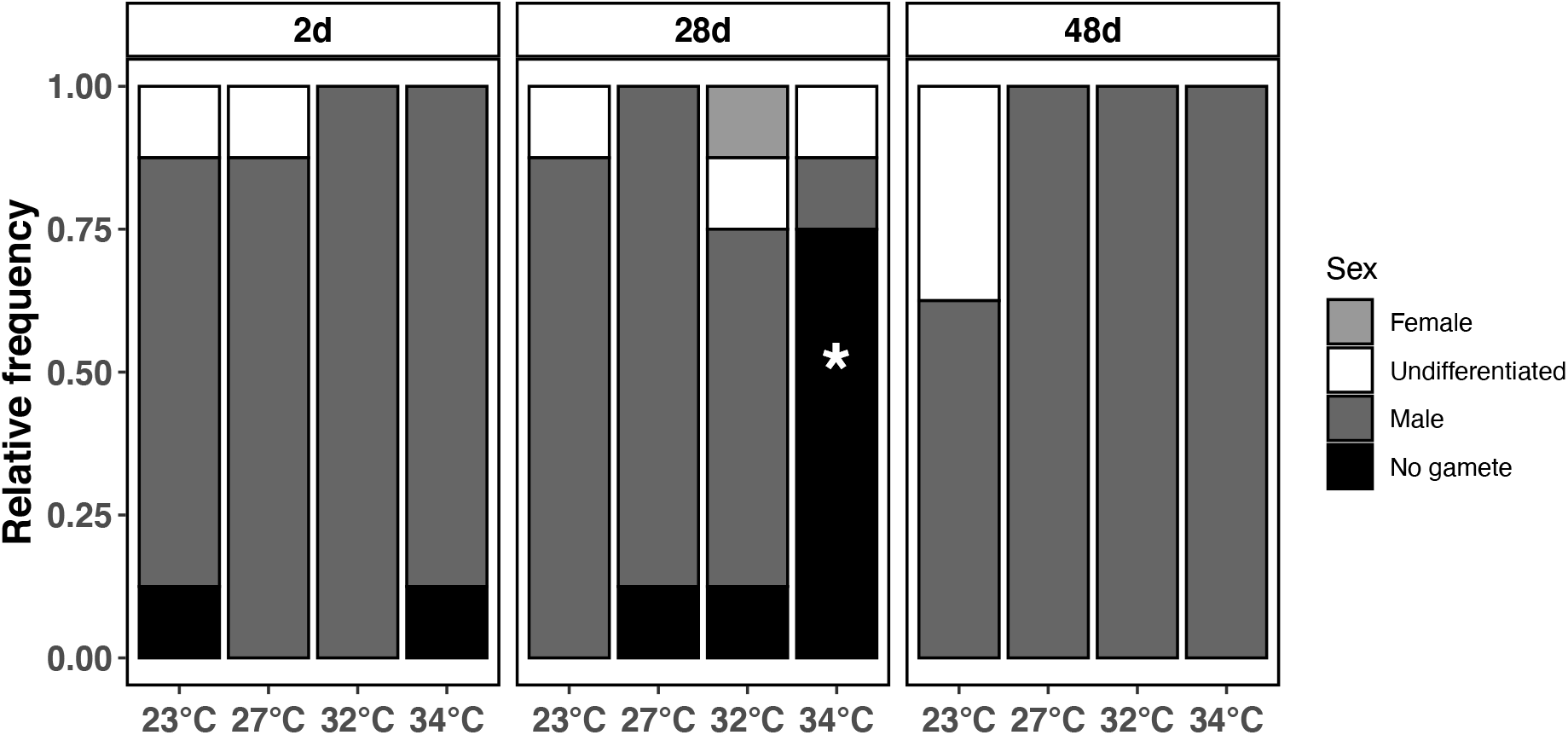
Categories of sex determination across sampling dates. Gametes were evaluated by gonads biopsy and microscopic examination. The asterisk represents the category mostly contributing to the difference across temperature treatments on day 28.

### (b) Genetic structuring, differentiation and candidate SNPs

We removed 6 individuals from the analysis that showed more than 15% missing genotypes, and identified 27,394 SNPs genotyped in 66 individuals (30 individuals from Gambier and 36 individuals from the Marquesas). The PCA showed the presence of two clearly separated clusters, matching the geographic/family origin of the samples (Fig. 2A). The importance of the geographical origin was further demonstrated by the high F_ST_ value of 0.21 (Supplementary Table S1, Fig. 2A). The contribution of each SNP to the variance of the DAPC is shown in Fig. 2B. In total, 263 SNPs were found in the top 1 percentile, and contribute the most to the discrimination between the two populations. These SNPs belonged to 161 transcripts, which showed an enrichment for GO categories including respiratory chain (cellular component, CC); NADH dehydrogenase (ubiquinone) activity (MF); NADH dehydrogenase activity (MF); regulation of alternative mRNA splicing, via spliceosome (Biological process, BP) and oxidoreductase activity (MF). A complete list of GO enrichment is provided in Supplementary Table S3.

**Fig. 2:**
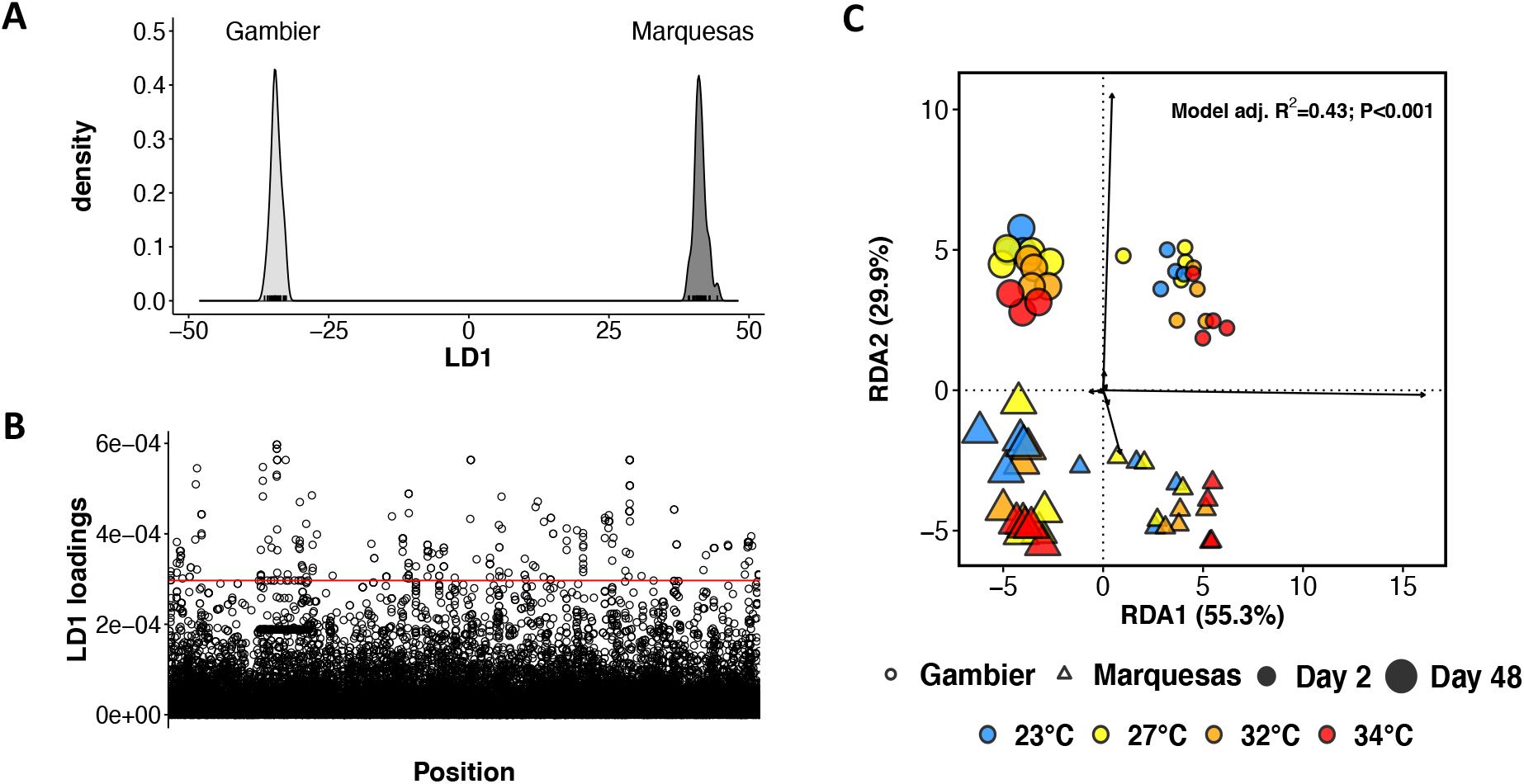
Genetic and genes expression variation across sampling dates, populations and temperature conditions. **A.** Clustering of populations through Discriminant Analysis of Principal Components (DAPC) based on SNPs variation (n = 27,394 SNPs) on the 1^st^ discriminant axis (LD1); **B.** Loadings of SNPs segregating population of the LD1. Red line indicates the top 1 percentile; **C.** RDA of gene expression level measured when the set temperature was reached (2 d) and at the end of the experiment (48 d).

### (c) Fatty acids

Temperature drove the changes in fatty acid (FA) composition of samples. After 48 days of exposure, the FA profile of oysters was strongly affected by temperature (RDA; ANOVA; *P* < 0.01; Fig. 3A), accounting for 33.49% of the total variance. Unsaturation index (UI) was negatively correlated with temperature (Pearson’s correlation; R = −0.87; *P* < 0.001; Fig. 3B), which was mainly explained by a reduced contribution of polyunsaturated fatty acids (PUFAs), like docosahexaenoic acid (DHA, 22:6n-3) and eicosapentaenoic acid (EPA, 20:5n-3, ANOVA; DHA, F = 31.95, *P* < 0.001; EPA, F = 20.17, *P* < 0.001), detected at higher temperatures. In contrast to these two PUFAs, arachidonic acid (ARA, 20:4n-6) increased with temperature (ANOVA; F = 7.77, *P* < 0.001). The FA profile of oysters was not affected by population (partial db-RDA; ANOVA; F = 2.31; *P* = 0.07). A full details of FA levels contents is provided in the Supplementary Table S1 and Supplementary Fig. S2).

**Fig. 3:**
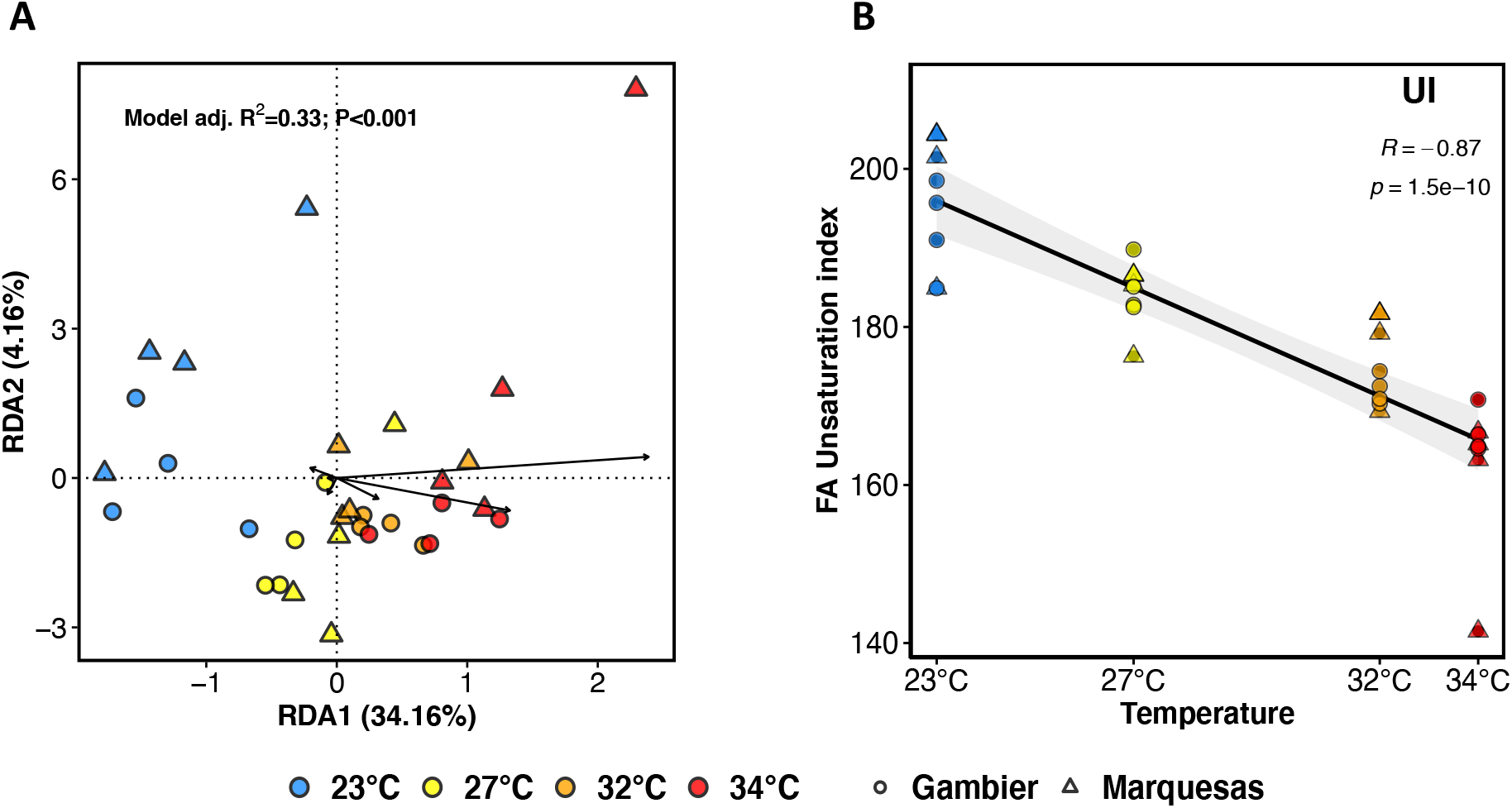
Fatty acid composition across temperature conditions and populations on day 48. **A.** Redundant discriminant analysis (RDA) of fatty acid profile (% FA) across populations and temperature treatment. **B.** Pearson’s correlation of unsaturated index.

### (d) Gene expression

The RDA model that included time, temperature and population explained 43.8% of the total variance in gene expression (RDA; ANOVA; *P* < 0.001). Time alone accounted for 25.9% of the variance explained by the model (ANOVA; partial db-RDA; F = 24.51; *P* < 0.001; Fig. 2C). Gene expression was also affected by population (ANOVA; partial db-RDA; F = 9.45; *P* < 0.001; Fig. 2C) and temperature (ANOVA; partial db-RDA; F = 7.14; *P* < 0.001; Fig. 2C), which explained 11.1% and 6.7% of the variance explained by the model, respectively.

### (e) Global genes plasticity in response to temperature

Overall, we found that more genes responded to temperature on day 2 (short-term stress response; N = 4,750) than after 48 days of exposure (acclimation response; N = 2,036; χ^2^ = 1224.9, *P* < 0.001; Table S1). Only 583 genes were affected by temperature at both 2 and 48 days of exposure. We modeled the reaction norms of temperature-responsive genes separately for each time and population. Mean *Tm* were only significantly different at day 2, with lower optimal temperature for genes showing U-shape pattern of expression in Gambier compared to Marquesas (Mann-Whitney U test; W = 50981; *P* < 0.001; Supplementary Table S2).

During the short-term exposure (2d), the plastic response to temperature involved enrichment of functions associated with immune system processes (BP), regulation of cell death and proliferation (BP), mRNA processing (BP), regulation of gene expression and epigenetic (BP), oxidoreductase activity (MF), ncRNA metabolic processes (BP), organelle organisation (BP), carbohydrate derivative binding (MF), mitochondrion (CC), generation of precursor metabolites and energy (BP), responses to stress including the synthesis of heat shock proteins (BP), and heat shock protein binding (MF).

After the long-term exposure (48d), temperature affected functions involved in mRNA processing (BP) and responses to stress (BP). We also found specific enrichment for NADH dehydrogenase complex (CC) and inner-mitochondrial membrane protein complex (CC) terms. A complete list of GO enrichment for each gene set is provided in Supplementary Table S3.

### (f) Molecular mechanism of short and long-term responses to elevated temperature

To identify putative molecular shifts due to the stress of acclimation, we contrasted the extreme-warm (34°C) and reference (27°C) conditions between days 2 and 48 and their interaction (34°C-2d *vs*. 34°C-48d) using pairwise comparisons for each population separately (Fig. 3).

On day 2, contrasts (27°C *vs*. 34°C) showed common enrichment across populations for enzyme-regulation activity (MF), regulation of cell death (BP), regulation of nitrogen compounds and protein metabolic processes (BP), regulation of proteolysis (BP) and regulation of response to stress (BP). The response to stress included at least five common HSPs: HSP71, HSP70, HSP70 cognate (Supplementary Fig. S3). We also found that oxidoreductase activity (MF) was enriched for the Marquesas population only. A complete list of GO enrichments is provided in the Supplementary Table S3.

On day 48 (27°C *vs*. 34°C), fewer genes (n= 206 and 274, for Gambier and Marquesas, respectively; Fig. 4) were differentially expressed and no GO term was significantly enriched between the conditions 27°C and 34°C. Notably, HSPs were not constitutively expressed on day 48 for both populations. Almost all inducible HSPs had low or null expression on day 48 for the Marquesas population. For the Gambier population nevertheless, HSP70, Hsc70-2, HSP70A1 and HSP71 were still more expressed at 34°C compared to 27°C, but at significantly lower levels that at day 2 (Supplementary Fig. S3).

**Fig. 4:**
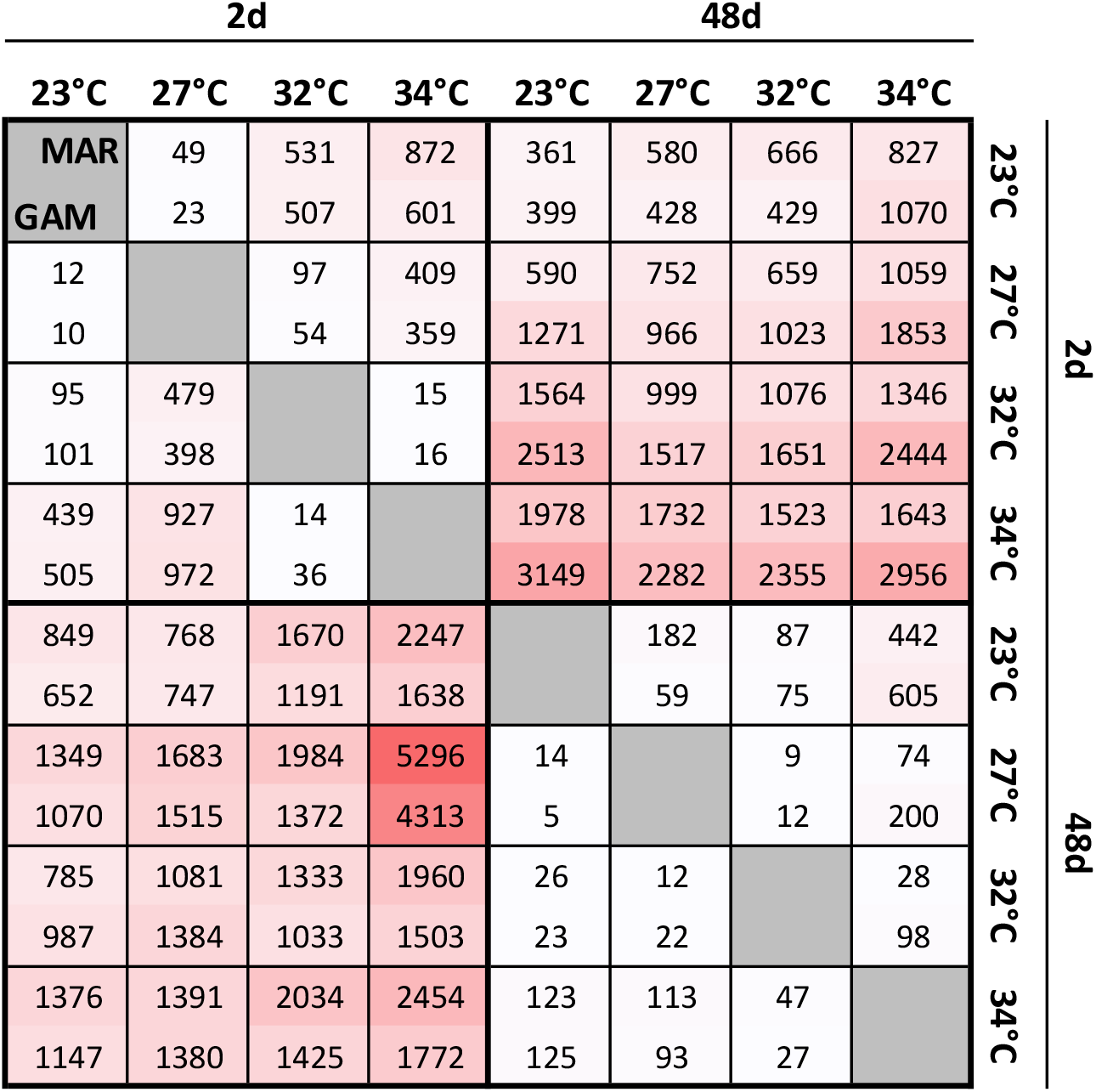
Number of responsive genes across time, populations and temperatures in pairwise contrasts. Above and below the diagonal (grey) are represented pairwise comparisons for the Marquesas (MAR) and Gambier (GAM) populations, respectively. Upper numbers represent genes up-regulated in treatment listed in the row; lower numbers represent genes up-regulated in treatment listed in the column. Genes response was considered significant when FDR < 0.01 and absolute fold-change (|FC|) > 2.

The molecular activity at 34°C varied across time. Up-regulated genes on day 2 show enrichment for responses to stimulus (BP) functions, regulation of lipid-metabolism processes (BF), carbohydrate derivative binding (MF), positive regulation of apoptotic process (BF), immune responses (BP), hydrolase activity (MF) and steroid hormone receptor activity (MF). In contrast, genes down-regulated at day 2 - 34°C compared to day 48 - 34°C, show enrichment for functions associated with the response and regulation of reactive oxygen species (ROS) in Gambier, but not in the Marquesas. However, both populations show enrichment for mitochondrial respiratory chain complex I (CC), peptide biosynthetic process (BP) and chemical homeostasis (BP).

### (g) Populations effect on gene expression

Populations had a profound effect on the overall gene expression (Fig.s 1D, 2). We found a total of 2,085 and 2,113 genes differentially expressed between populations on days 2 and 48 of exposure, respectively. The population effect on day 2 included differences in DNA polymerase activity (MF), rRNA binding (MF), peptide and amide synthesis processes (BP), cellular nitrogen compound metabolic processes (BP), ribonucleoprotein complex (CC) and oxidoreductase activity (MF). We found enrichment for oxidoreductase activity (MF) and cellular nitrogen compound metabolic processes (BP), which is independent of time. On day 48 we also detected enrichment of sulfuric ester hydrolase activity (MF). Interestingly, we found overlap across candidate genes identified as outlier functional SNPs (non-synonymous mutation) and genes differentially expressed between populations. This overlap (n = 14 transcripts) included central actors in mitochondria functioning such as mitochondrial amidoxime-reducing component 1 (MARC1) and ATP synthase subunit O, mitochondrial (ATPO) coding genes and two transcripts coding for the matrix protein N66, a regulator of shell calcification in molluscs (Rivera-Perez, Ojeda-Ramirez de Areyano, & Hernandez-Saavedra, 2019).

Finally, relatively few genes (17 and 10 on days 2 and 48, respectively) showed a significant pattern of genotype-by-environment interaction (GEI), with no significant (Bonferroni *adj. P* < 0.1) enrichment detected.

## Discussion

Ocean warming is a particularly challenging threat for tropical marine bivalves species because most of them live already near their upper thermal limits (Huey et al., 2009; Stuart-Smith, Edgar, & Bates, 2017). The thermal sensitivity of organisms is a strong contributor to the biogeographic boundaries of populations and species (Hochachka & Somero, 2002). Here, we show that tropical sessile organisms might be able to cope with abnormally elevated temperature on long-term (several weeks) and that divergent populations, naturally experiencing contrasted environmental conditions, show genetic variation associated with central actor of the response to heat stress.

### (a) Acclimation potential to long-term exposure to elevated temperatures in Pinctada

The detrimental effect of elevated temperature was partially compensated on the long-term, revealing an unsuspected acclimation potential in *P. margaritifera*. We show that exposure to elevated temperature induces a rapid activation of molecular indicators of systemic stress (*e.g* HSP) and is accompanied by a significant reduction in gamete production (75% of the individuals showing no gametes) at 28 days, all together strongly supporting the notion that 34°C lays beyond the species’ thermal range (Kooijman, 2000; Sokolova, 2013; Sokolova, Frederich, Bagwe, Lannig, & Sukhotin, 2012). Our results support previous observations that estimated the optimal temperature range of *P. margaritifera* to fall between 23°C and 29°C, depending on the source population, and negative SFG after seven days of exposure to 34°C (Le Moullac et al., 2016; Yukihira, Lucas, & Klumpp, 2000). However, after a longer exposure (48 days) to 34°C, we show that gonads were differentiating again. Gamete production indicates that sufficient energy was available for basal somatic maintenance and to supply at least part of the reproduction (Sokolova, 2013). The discrepancy between the short- and long-term observations reveals an overall physiological capacity of *P. margaritifera* to adjust to stressful thermal conditions. Similarly, the unsaturation index show a steady decrease, as predicted by the homeoviscous adaptation theory (Sinensky, 1974), with no signs of plateau suggesting that individuals are well acclimated at 34°C after 48 days exposure. These observations remain true for the time and conditions of the experiment (steady temperature control, feeding *ad libitum*, continuous oxygen supply and quality-controlled water) and might not imply the resilience of *Pinctada* populations when facing a combination of multiple stressors (acidification, oxygen deprivation, etc.). Furthermore, study in marine ectotherms showed that thermal tolerance varies across life stages (increased from embryos to adults) (Dahlke et al., 2020), hence, basing evidence only from adults’ response might underestimate the effect of global warming on the species’ resilience. Nevertheless, these capacities have so far been largely underestimated, especially with the over-representation of short-term experiments (Semsar-kazerouni & Verberk, 2018); consequently, the associated physiological and molecular mechanisms remain largely unknown.

### (b) Mechanisms of acclimation to long-term extreme temperatures exposure

The HSR was activated after 2 days exposure to 34°C and decreased thereafter. The HSP turnover is energetically demanding, representing up to 10% of the total protein synthesis costs (Hofmann & Somero, 1995; Sokolova, 2013). Therefore, covering both maintenance and reproduction costs under prolonged exposure to extreme temperatures, as we show here, would necessarily require other physiological adjustments, different from the highly energy demanding HSR. The long-term acclimation (between 2 and 48 days) is characterized by an active solicitation of the activity of detoxification and mitochondrial machinery, notably via the respiratory chain complex I. Besides its role on regulating apoptosis and ROS, the complex I catalyzes the entry of electrons into the mitochondria; thus, any functional change might be tightly correlated with physiological limits of the global respiration machinery (Sharma, Lu, & Bai, 2009). The activation of respiration machinery is in line with the oxygen-limitation hypothesis that drives the thermal tolerance of organisms (Dahlke et al., 2020; Pörtner, 2010).

In parallel, we showed a strong remodelling of membrane fatty acids that is consistent with the HVA theory (Sinensky, 1974). Maintenance of membrane fluidity is an essential component of the acclimation process and also mediate respiration machinery (Budin et al., 2018; Hazel, 1995; Fabrice Pernet et al., 2007). Our observations agree well with previous data on marine temperate mollusc species exposed to elevated temperatures, which showed decreases in 22:6n-3 and 20:5n-3 with increasing temperatures (Muir, Nunes, Dubois, & Pernet, 2016; Pernet et al., 2007). Therefore, lipids remodelling, specially PUFAs, is also a common mechanism of thermal acclimation in marine ectotherms including stenothermal tropical species.

### (c) Evidence for adaptive divergence between populations

Gambier and Marquesas populations are phenotypically (shell morphology and coloration) and genetically different (Reisser et al., 2019), and this is maintained under laboratory conditions in the F1 progeny. Gene expression varied markedly between populations but not membrane lipids. In previous studies, we found differences in remodelling of membrane lipids among populations of bivalves. For example, the pattern of lipid remodelling in response to temperature in oysters *C. virginica* and clams *Mercenaria mercenaria* shows strong intraspecific variation despite the fact that animals were maintained in the same environment (Parent, Pernet, Tremblay, Sévigny, & Ouellette, 2008; Pernet, Tremblay, Gionet, & Landry, 2006; Pernet, Tremblay, Redjah, Sevigny, & Gionet, 2008). In these studies, however, intra-specific variations in lipid remodeling were associated with differences in metabolic rates and growth rates. Here we did not observe any differences in growth or gamete production between populations and this could explain similarities in FA profiles.

Here we show that a large part of the gene expression variation was significantly controlled by the genotype, with a limited GEI effect, and that difference was pervasive after almost three months exposure to common environment (acclimatization in the wild and in laboratory). While population (heritable) effects are often confounded with carry-over effects (life-history, family or early life stages) (Kellermann et al., 2017), the physiological condition at the beginning of the experiment did not differ among populations, strongly suggesting that their responses were most likely genetically-coded.. Among the genes showing both genetic and expression-level variation, we identified the N66 matrix protein-coding gene, a major actor of the shell biomineralization in pearl oysters (Rivera-Perez et al., 2019). In parallel, biological function enrichment analyses showed that oxidoreductase activity and cellular nitrogen compound metabolic pathways depended constitutively on the population. Similarly, non-synonymous genetic variation between populations also revealed differences in oxidoreductase activity, mainly through NADH dehydrogenase activity, a complex involved in the mitochondrial oxidative phosphorylation system (OXPHOS). Mutation on the mitogenome, particularly the OXPHOS genes complex, can have a profound effect on respiration and overall metabolism in animals, thus playing a key role in their adaptation to climate change (Lamb et al., 2018; Sokolova, 2018). Similarly, variation in the expression of complex I genes (including NADH:ubiquinone oxidoreductase) differentiated ecologically divergent populations of temperate fishes (and F1 progenies) (Narum & Campbell, 2015). We interpret the data as a mark of adaptive divergence across the Marquesas and Gambier population systems, although further studies including multiple populations’ genetic surveys are warranted to test this hypothesis.

## Conclusions

Understanding and quantifying the response of organisms and populations to ongoing and projected global changes is critical for environmental-resource managers and policy makers. Thermal stress is undoubtedly one of the most concerning threats to marine tropical ectotherms. We showed that *P. margaritifera* has acclimation capacities to elevated temperatures largely underestimated to date. Individual- and molecular-level responses showed major differences across conditions and time. The time component is often ignored in short-term experiments and can mislead predictions of long-term dynamics. Finally, while the molecular response to temperature was largely shared between populations, there was a marked divergence for loci involved in the respiration machinery. We now urge for comprehensive studies integrating long-term monitoring (several developmental stages and/or several generation) that would integrate populations diversity to draw significant patterns and accurate predictions.

## Ethics

Specimens used in this study were collected and held under a special permit delivered from the French Polynesian government.

## Data accessibility

Raw sequencing RNAseq data have been made publicly available on NCBI portal (PRJNAXXXXX).

## Authors’ contributions

FP and JLL conceived the experiment. JLL and CS conducted the experiment. CLK provided the biological material through its partnership network. CB carried out the laboratory benchwork for the transcriptomic data. FP conducted lipid analyses. CJM, CR, FP, GLM, JLL and LM analyzed the data. CJM, CR, JLL and FP wrote the manuscript. All co-authors contributed substantially to reviewed drafts of the manuscript.

## Competing interests

We declare that we have no competing interests.

## Funding

This work was supported by joint grant from Labex MER-CORAIL (project Gamma) and funding for wild stock collection in Marquesas archipelago through the AmeliGEN project (#10065/MEI/DRMM) supported by grant from the “Direction des Ressources Marines” of French Polynesia.

## Acknowledgements

We are grateful to Seiji Nakasai and Dominique Devaux for conducting animal breeding, rearing and providing F1 individuals from their hatchery located in Gambier archipelago. We also thank Manaarii Sham Koua for his precious support in conducting animal breeding and rearing for the F1 Marquesas individuals at the Ifremer’s research center (Tahiti, French Polynesia). We also thank Laurianne Bish for continuously providing the microalgae material and Claudie Quéré and Valérian Leroy for conducting lipid analyses. We finally thanks Vaihiti Teaniniuraitemoana and Virgile Quillien for their help during sampling.

## Notes

### Competing Interest Statement

The authors have declared no competing interest.

